# A parent-of-origin effect on embryonic telomere elongation determines telomere length inheritance

**DOI:** 10.1101/2025.01.28.635226

**Authors:** Hyuk-Joon Jeon, Mia T. Levine, Michael A. Lampson

**Affiliations:** Department of Biology, School of Arts and Sciences, University of Pennsylvania; Philadelphia, PA 19104, USA; Penn Center for Genome Integrity, University of Pennsylvania; Philadelphia, PA, USA

## Abstract

Telomere length is inherited directly as a DNA sequence and as a classic quantitative trait controlled by many genes across the genome. Here, we show that neither paradigm fully accounts for telomere length inheritance, which also depends on a parent-of-origin effect on telomere elongation in the early embryo. By reciprocally crossing mouse strains with different telomere lengths, we find that telomeres elongate in hybrid embryos only when maternal telomeres are short and paternal telomeres are long. In the reciprocal cross, telomeres shorten. These differences in embryonic telomere elongation, which emerge before zygotic genome activation, predict adult telomere length. Moreover, when telomeres do elongate, we find molecular signatures of a recombination-based mechanism of telomere elongation, called the Alternative Lengthening of Telomeres (ALT) pathway, previously suggested to elongate telomeres in the pre-implantation embryo. We propose that ALT is triggered by a combination of genetic asymmetry in telomere length and epigenetic asymmetry between maternal and paternal chromosomes in the zygote. Our findings offer new insight into the complex interaction of genetic and epigenetic determinants of telomere length inheritance.

## Introduction

Telomeres are composed of specialized DNA, RNA, and proteins that interact to preserve the integrity of chromosome termini.^1,2^ Despite this vital function, telomere length varies dramatically across human populations; that is, copy number of the telomeric satellite, TTAGGG, is highly variable. For example, within sub-Saharan Africa alone, average leukocyte telomere length ranges from 6kb to 11kb.^3^ Over the past two decades, genome-wide association studies have identified multiple genes that explain a portion of the quantitative variation in telomere length.^4^ Pathogenic mutations in some of these genes cause extremely long or extremely short telomeres. Both extremes are associated with multiple cancers.^5^ These validated genes encode proteins that span the telomerase holoenzyme, the telomere protection complex, and DNA repair factors. Such genes mediate telomerase access to chromosome ends and exposure of ends to nucleolytic degradation. These discoveries support the intuitive idea that telomere length is a classic quantitative trait controlled by many genes acting *in trans* to promote telomere integrity. Inheritance of telomere length should, therefore, depend on the alleles of these genes inherited from the parents.

Unlike other polygenic traits, however, telomere length is also a trait itself, transmitted directly from germ cell to zygote.^6^ Offspring, at least initially, inherit their parents’ telomeric satellite copy numbers. Supporting this direct inheritance paradigm, analysis of human telomeres suggests that telomere length in the gametes explains variation in adult offspring independent of allelic variation at genes.^7^ In further support of this paradigm, intercrossing short-telomere mice that are heterozygous mutant for a telomerase component (*Tert^+/−^)* yields wildtype (*Tert^+/+^*) progeny that maintain the same short telomeres for multiple generations despite the presence of a functional telomerase.^8^ These human and mouse data are consistent with direct inheritance of telomere length, independent of genotype.

Empirical data from trios introduces a third paradigm. Studies of various vertebrate species have documented parent-of-origin effects, where the offspring telomere length correlates with either the paternal or maternal telomere length (reviewed in^6^). Moreover, human paternal age effects linked to longer sperm telomeres in older fathers have been widely reported.^9–14^ Parent-of-origin effects are difficult to reconcile with the polygenic trait paradigm and suggest a potentially complex contribution of direct inheritance.^15^ Unraveling this complexity has been particularly challenging given the confounding effects of the environment, age, cell type, and genetic background on telomere length inheritance in humans. Currently, a mechanism to explain maternal or paternal effects on telomere length is lacking.

We set out to establish a controlled, experimental model to evaluate the alternative predictions from the three paradigms (Figure 1A). Specifically, we crossed closely related species or inbred strains of mice with different telomere lengths and measured F_1_ progeny telomere length by fluorescence *in situ* hybridization (FISH). If telomere length is inherited directly as a DNA sequence, F_1_ telomeres should have an average telomere length that is intermediate relative to the parents and high variance due to a bimodal distribution. If telomere length is inherited as a polygenic trait, then F_1_ telomeres should also have average telomere length that is intermediate relative to the parents but with low variance. Finally, a parent-of-origin effect predicts F_1_ telomeres should have an average telomere length that is similar to one parent only and have low variance. Using our mouse model system, we find evidence of a parent-of-origin effect on telomere elongation in the early embryonic cell cycles, leading to an apparent paternal effect on telomere length. Elongation correlates with molecular signatures of the recombination-based alternative lengthening of telomeres (ALT) pathway. Our findings suggest that a combination of directly inherited maternal-paternal genetic asymmetry and epigenetic differences between parental chromosomes in the embryo determine telomere inheritance.

**Figure 1.**
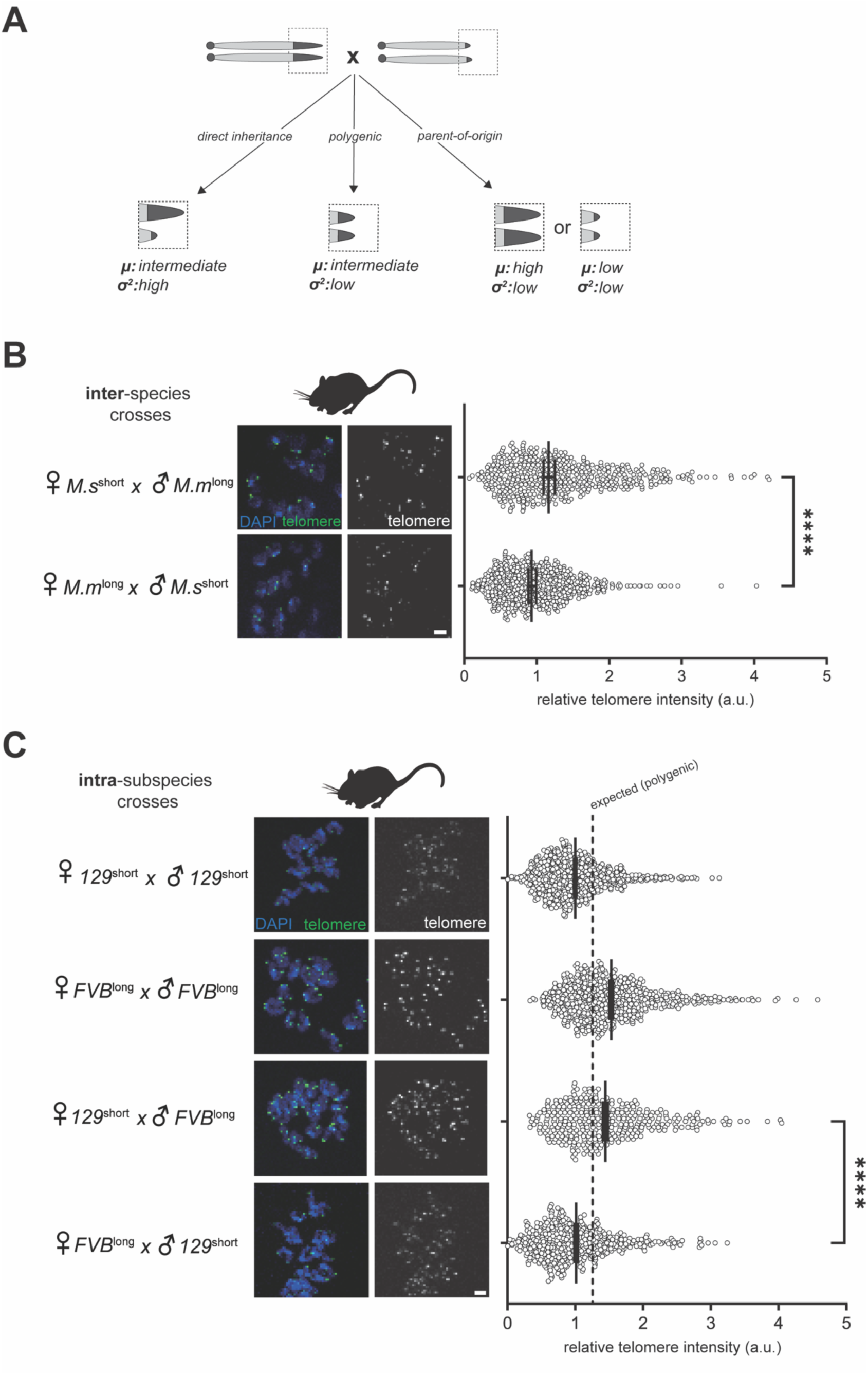
Parent-of-origin effects on telomere length. (**A**) The three paradigms of telomere length inheritance make different predictions for mean (μ) and variance (σ^2^) of progeny telomere length from crosses between parents with divergent telomere lengths. Direct inheritance (left) predicts that parental telomeres are maintained in the F_1_ progeny; consequently, offspring telomere length should be intermediate but with high variance between telomeres. The polygenic trait paradigm (middle) predicts intermediate telomere length with low variance. A parent-of-origin effect (right) predicts that telomeres in the progeny match one of the parents and have low variance. (**B, C**) Telomere FISH of metaphase II eggs from hybrid adult progeny of reciprocal crosses between *Mus musculus* (FVB strain) and *Mus spretus* (B), or between *Mus musculus domesticus* strains 129^short^ and FVB^long^ (C). Data from pure 129^short^ or FVB^long^ eggs are also shown in (C). Representative images are shown; scale bar 5 µm. Graphs show intensities measured for individual telomeres. For each sample, n ≥ 375 telomeres of individual chromatids were measured from at least 86 eggs from at least 3 females pooled from 2 independent experiments for (B), or n ≥ 363 telomeres of chromatids from at least 51eggs from at least 6 females pooled from 2 independent experiments for (C). For each experiment, data were normalized to the mean of the *Mus musculus*♀ x *Mus spretus*♂ cross (B) or to the 129^short^ mean (C). Bars: mean and standard error of the mean. \*\*\*\**p* < 0.0001.

## Results

### Parent-of-origin effects on telomere length

Determining the inheritance pattern of any trait requires variation. To study telomere length inheritance, we selected mouse species and strains known to have dramatic differences in telomere length. *Mus spretus* has short telomeres, while closely related, standard laboratory strains of its sister species, *Mus musculus,* generally have longer telomeres.^16^ However, short telomere laboratory strains of *Mus musculus* have also been described.^17^ We used reciprocal crosses either between *Mus musculus* and *Mus spretus*, or between selected *Mus musculus* strains with different telomere lengths, to investigate the determinants of telomere length inheritance.

The two classes of progeny that arise from reciprocal crosses are genotypically identical and so differ only in the parent-of-origin of the long (or short) telomere. Our assays of the F_1_ adult progeny focused on their gametes because the gamete represents both the focal adult offspring and the telomere length that would be transmitted to the following (F_2_) generation. To measure telomere length, we hybridized a FISH probe cognate to the telomeric repeat in adult female gametes (we find minimal differences between male and female gametes, Figure S1).

Reciprocal crosses between *Mus musculus* (long telomeres) and *Mus spretus* (short telomeres) revealed that progeny telomere length depends on the direction of the cross. Adult offspring matched the father, with longer telomeres when their fathers were the long-telomere *M. musculus* (Figure 1B) or shorter telomeres when their fathers were the short-telomere *M. spretus*. This interspecies cross rejects both the polygenic trait and simple direct inheritance paradigms and instead implicates a parent-of-origin effect.

To conduct the same experiment with parents who are more genotypically similar but still differ in telomere length, we identified from the literature two *M. musculus domesticus* strains: a long-telomere strain (FVB) and a short-telomere strain (129).^17^ Our single telomere FISH measurements in adult gametes confirmed the previous report from adult liver: the 129 strain has short telomeres relative to the FVB strain (Figure 1C, 129^short^ and FVB^long^ hereafter). Reciprocally crossing the long- and short-telomere *Mus musculus* parents revealed that adult F_1_ telomere length depends on the direction of the cross (Figure 1C). F_1_ telomeres from the FVB^long^♀ x 129^short^♂cross were short. Conversely, F_1_ telomeres from the 129^short^♀ x FVB^long^♂ cross were long. This paternal effect recapitulates the between-species reciprocal cross (Figure 1B) and demonstrates that neither the direct inheritance nor the polygenic trait paradigms (Figure 1A) fully account for telomere length inheritance.

### Parent-of-origin effects on telomere elongation during preimplantation development

Telomerase activity in stem cells maintains telomere length, counteracting both the end- replication problem and oxidative stress that degrade telomere ends.^18^ During pre-implantation development, telomeres have been reported to elongate rather than simply maintain length, setting an initial telomere length for the developing organism.^19,20^ These observations motivated us to probe the source of the paternal effect in the embryo, leveraging the experimentally tractable reciprocal crosses between *Mus musculus domesticus* inbred strains.

We first analyzed telomere length in blastocysts, the last stage of preimplantation development. The relative telomere lengths from reciprocal crosses between the two different strains recapitulated the observations in adult progeny (Figure 1C), with blastocyst telomere length dependent on the direction of the cross (Figure 2A). Telomeres were relatively long in the 129^short^♀ x FVB^long^♂ cross, or relatively short in the reciprocal cross. These findings suggest that differences in telomere elongation during preimplantation development explain the observed paternal effect on telomere length inheritance in adults.

**Figure 2.**
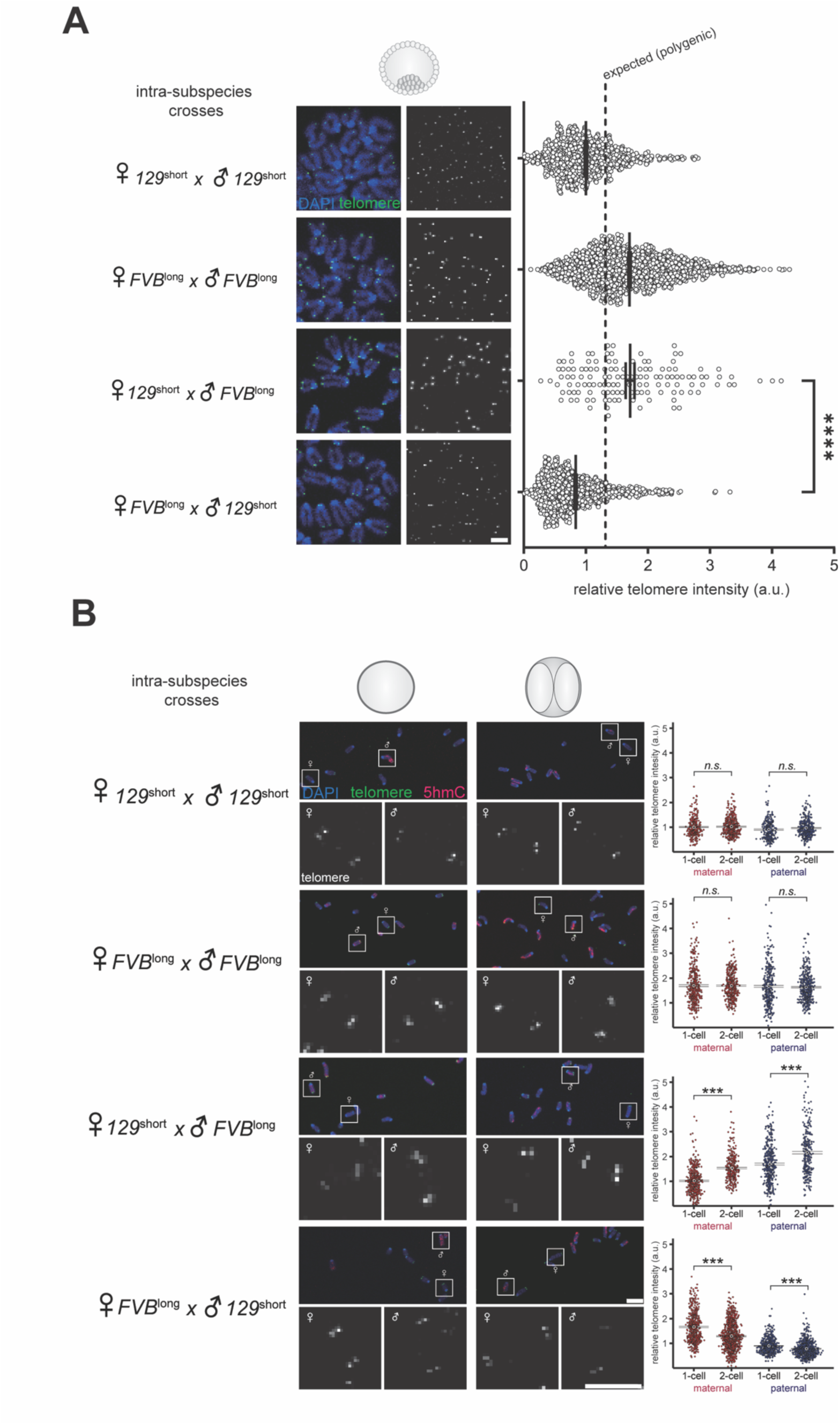
Parent-of-origin effects on telomere elongation during preimplantation development. (**A**) Telomere FISH of embryos at the morula to blastocyst stage from pure 129^short^ or FVB^long^ strains or from reciprocal crosses between the two strains. Representative images are shown, and graphs show intensities measured for individual telomeres. For each sample, n ≥ 133 telomeres of chromatids were measured in at least 30 eggs from at least 4 females pooled from at least 2 independent experiments. Dotted vertical line indicates the intermediate telomere length expected from the polygenic inheritance paradigm. We note that development *in vitro* of the ♀ 129^short^ x ♂ FVB^long^ was less efficient compared to other crosses, resulting in fewer blastocysts. For each experiment, data were normalized to the ♀ 129^short^ x ♂ 129^short^ embryo mean. Bars: mean and standard error of the mean (**B**) Measurements of telomere elongation between the first and second mitoses in embryos from pure 129^short^ or FVB^long^ strains or reciprocal crosses between the two strains. One- and two-cell embryos were fixed simultaneously and processed for telomere FISH and immunostaining for 5hmC to label paternal chromosomes. Representative images are shown, and graphs show intensities measured for individual telomeres. For each sample, n ≥ 1291 paired telomeres of sister chromatids were measured from at least 112 embryos from at least 8 females pooled from at least 2 independent experiments. All data were normalized to the mean of maternal telomeres in one-cell ♀ 129^short^ x ♂ 129^long^ embryos. Bars: mean and standard error of the mean. Scale bars, 5 µm. \*\*\*\**p* < 0.0001.

To determine the source of this variation between genotypically identical progeny, we focused on elongation between the first and second embryonic mitoses when the maternal and paternal direct inheritance should have an outsized impact relative to the previously silent zygotic genome. Given the observed paternal effect, we measured relative telomere lengths of maternal and paternal chromosomes separately (Figure 2B). To distinguish maternal chromosomes from paternal chromosomes, we took advantage of the intermediate methylation state (5- hydroxymethylcytosine, or 5hmC) that marks paternal chromosomes immediately following fertilization. Maternal chromosomes, in contrast, are methylated (5mc).^21^ We find that telomere length neither elongates nor degrades in embryos from inbred strains. In embryos from the 129^short^♀ x FVB^long^♂ cross, both sets of parental telomeres elongate, consistent with the relatively long telomeres observed at the blastocyst stage. In contrast, maternal telomeres shorten in the reciprocal FVB^long^♀ x 129^short^♂ cross, consistent with the relatively short telomeres in blastocysts. These findings indicate that telomere length inheritance depends on regulation during the earliest cell cycles in the embryo, ostensibly independent of the products of zygotic genome activation at the two-cell stage. This parent-of-origin effect on elongation explains the paternal effect on telomere length.

### Parent-of-origin effects on ALT during preimplantation development

A classic mechanism underlying parent-of-origin effects is maternal deposition of protein or RNA into the embryo. For telomere length regulation, these maternally provisioned products might be telomerase components or DNA repair factors. However, this mechanism alone fails to account for the observed parent-of-origin effect on telomere elongation: despite the same maternal provisioning, embryos from 129^short^ mothers elongate when the paternal telomeres are long (129^short^♀ x FVB^long^♂) but not when the paternal telomeres are short (129^short^♀ x 129^short^♂) (Figure 2B). Moreover, previous findings suggest that mouse telomeres elongate in the absence of telomerase activity^19^, instead using a recombination-based mechanism called Alternative Lengthening of Telomeres (ALT).^22^ We hypothesized that the parent-of-origin effect on telomere elongation is due to differences in ALT activity that depend on the relative lengths of the maternal and paternal telomeres.

Telomere invasion for homologous recombination begins with the formation of 3’ (G-rich) and, possibly, 5’ (C-rich) telomeric overhangs.^23,24^ ALT activation in cancer cells is typically assessed by the presence of single-stranded telomeric DNA or by markers for DNA damage at telomeres, which can generate 3’ overhangs ^25–27^. If this ALT-initiating event determines embryonic telomere elongation, we would expect comparatively more such overhangs in the 129^short^♀ x FVB^long^♂ embryos that elongate telomeres. Given prior studies in cancer cells indicating that ALT is active in S-G2 phase of the cell cycle^28,29^, we probed for the ALT-initiating single-stranded telomeric overhangs in G1 of two-cell embryos (Figure 3A). We labeled G-rich telomeric sequences by FISH under non-denaturing conditions (Native-FISH), as these 3’ overhangs would result from the end replication problem in the first S-phase. We find significantly more G-rich, single-stranded telomeric foci in embryos from 129^short^ mothers compared to embryos from FVB^long^ mothers, regardless of the identity of the father (Figure 3B). This finding suggests that the presence of 3’ overhangs depends on both short telomere length and a maternal-specific chromatin state. Furthermore, the restriction of elongation only to 129^short^♀ x FVB^long^♂ embryos suggests that elevated overhangs are necessary but not sufficient to promote elongation. Elongation requires both 3’ overhangs and long telomeres as targets for invasion.

**Figure 3.**
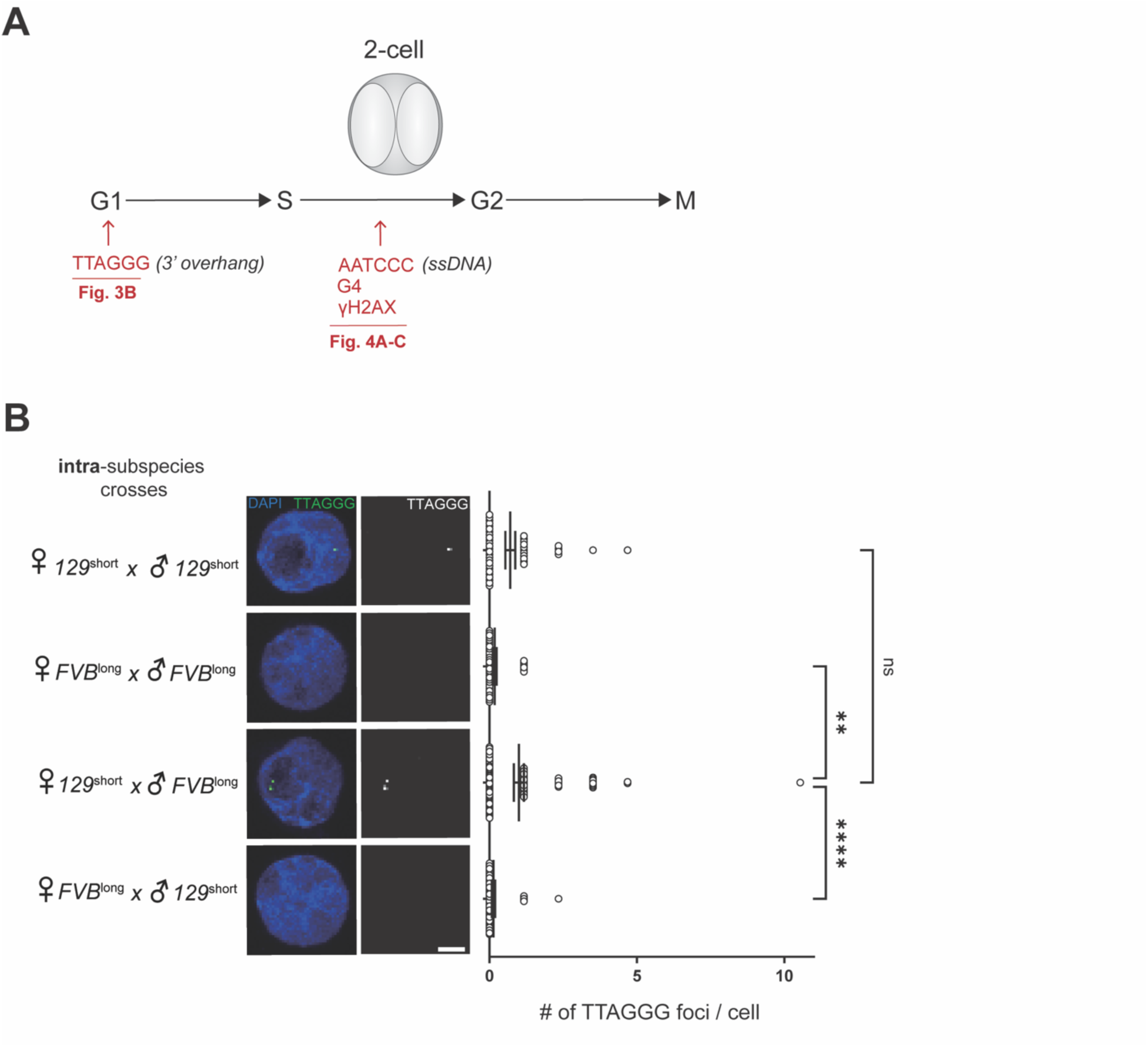
Parent-of-origin effects on the presence of G-rich single-stranded telomeric DNA in two-cell embryos. (**A**) Schematic of time points when signatures of ALT were assayed in two-cell embryos. Native- FISH was used at G1 to label 3’ single-strand overhangs (TTAGGG) and at G2/S to label 5’ single-strand overhangs (AATCCC). Immunofluorescence was used to label G4 structures and DNA damage signaling (γH2AX) at G2/S. (**B**) Native-FISH labeling 3’ G-rich single-stranded telomeric DNA in G1-stage two-cell embryos from pure 129^short^ or FVB^long^ strains or reciprocal crosses between the two strains. Representative images are shown; scale bar, 5 µm. Graph shows the number of foci per nucleus (each dot represents one nucleus). For each sample, n ≥ 39 nuclei were measured from at least 23 two-cell embryos from at least 4 females pooled from at least 2 independent experiments. Bars: mean and standard error of the mean. \*\*\*\**p* < 0.0001.

To test for ALT in S-G2 of two-cell embryos, we examined three established readouts. First, we used Native-FISH to label C-rich, single-stranded telomeric DNA, which is a specific marker for ALT during or after S-phase.^30^ Second, we stained for G-quadruplex (G4) structures promoted by the presence of single-stranded, telomeric G-rich DNA and known to be associated with DNA damage and ALT, together with the telomere-binding protein TRF1 to label telomeres. Third, we stained for γH2AX as a marker for DNA damage, together with TRF1. We find that all three ALT markers are elevated in the elongating 129^short^♀ x FVB^long^♂ embryos compared to the other three crosses: in only this cross did we observe elevated numbers of foci with C-rich telomeric sequences (Figure 4A) and elevated signals of G4 structures and DNA damage overlapping with telomeres (Figure 4B and C, respectively). We also note that embryos from 129^short^ females have more DNA damage overall than embryos from FVB^long^ females, regardless of the male parent, suggesting a strain-specific maternal effect on DNA damage (Figure S2). However, telomeres elongate in 129^short^♀ x FVB^long^♂ embryos but not in 129^short^♀ x 129^short^♂embryos (Figure 4C), suggesting that elevated overall damage is insufficient to trigger elongation. Together, these data demonstrate that ALT readouts in S-G2 are elevated only in embryos that elongate telomeres among the four cross-types (Figure 2B).

**Figure 4.**
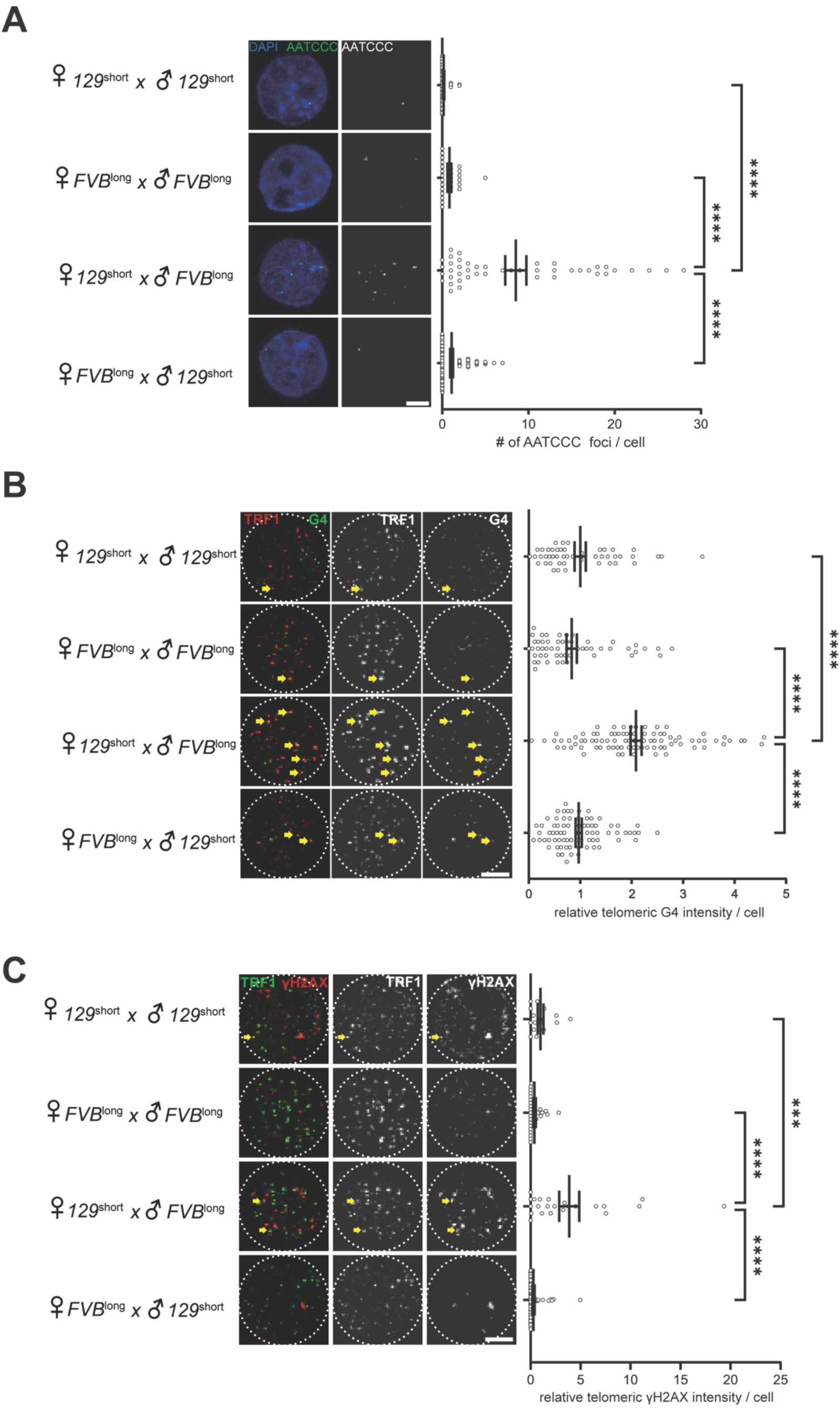
Parent-of-origin effects on ALT in two-cell embryos. Two-cell embryos from pure 129^short^ or FVB^long^ strains or reciprocal crosses between the two strains, were fixed in S-G2 and processed for Native FISH or immunostaining. (**A**) Native-FISH labeling C-rich single-stranded telomeric DNA. Representative images are shown, and graph shows the number of foci per nucleus (each dot represents one nucleus). For each sample, n ≥ 26 nuclei were measured from at least 22 embryos from at least 5 females pooled from at least 2 independent experiments. Bars: mean and standard error of the mean. (**B, C**) Immunostaining for G4 structures (B) or γH2AX (C), together with TRF1. Representative images are shown, with yellow arrows indicating telomere regions labeled with TRF1. Graphs show γH2AX or G4 intensity colocalizing with TRF1. Each dot represents intensity summed over one nucleus. For each sample, n ≥ 47 nuclei were measured from at least 24 two-cell embryos from at least 5 females pooled from at least 2 independent experiments for G4, or n ≥ 16 nuclei were measured from at least 10 two-cell embryos from at least 3 females pooled from at least 2 independent experiments for γH2AX. For each experiment, data were normalized to the mean of 129^short^♀ x 129^short^♂ embryos. Bars: mean and standard error of the mean. Scale bars, 5 µm. \*\*\**p* < 0.001, \*\*\*\**p* < 0.0001.

## Discussion

Parent-of-origin effects classically implicate epigenetic mechanisms. For telomere length, parent-of-origin effects can also include the direct transmission of telomere satellite copy number. Our data suggest that when maternal and paternal telomere lengths are different, a complex interaction of both mechanisms impacts telomere length inheritance. This interaction determines telomere elongation dynamics in the early embryo and, ultimately, adult telomere length. Specifically, telomeres elongate when paternal telomeres are long and maternal telomeres are short, leading to paternal-like, long telomeres in the F_1_ adult. Conversely, telomeres shorten when paternal telomeres are short and maternal telomeres are long, leading to short telomeres in the F_1_ adult. Finally, telomere length remains constant when parental telomeres are symmetrical. These observations suggest that a parent-of-origin effect on telomere *elongation* governs the apparent paternal effect on telomere *length*. A previous study also used reciprocal crosses to study the effects of telomere length variation using telomerase null and wild-type parents. Long-telomere wildtype fathers crossed to short telomere Tert*^−/−^* mothers yielded almost 3x more viable morula stage embryos than the reciprocal cross^31^, consistent with a parent-of-origin effect on telomere regulation. In both cases, a parent-of-origin effect strongly implicates an epigenetic mechanism.

We propose a three-step model to explain the observed embryonic maternal and paternal telomere elongation in crosses between mothers with short telomeres and fathers with long telomeres (Figure 5). First, short maternal telomeres initiate ALT with the abundant G-rich 3’ overhangs available in G1 of two-cell embryos (Figure 3). We suspect that the additional 3’ overhangs detected in short telomere mothers arise from the challenge of forming a stable telomere loop (T-loop) when telomeres are short^32^, so that telomeres are dynamically transitioning between T-loop formation and single-stranded DNA. Given that we observed elongation only in the presence of long paternal telomeres, we propose that these paternal telomeres serve as targets for invasion by maternal telomeres and as templates for maternal telomere elongation. This process would generate single-stranded paternal telomeric DNA during elongation, which is known to adopt G4 structures that induce DNA damage, thus initiating ALT to elongate paternal telomeres. This model accounts for all of our observations in two-cell embryos: elevated single-stranded G-rich telomeric DNA in G1 and single-stranded C-rich telomere DNA, DNA damage, and G4 structures in S-G2.

**Figure 5.**
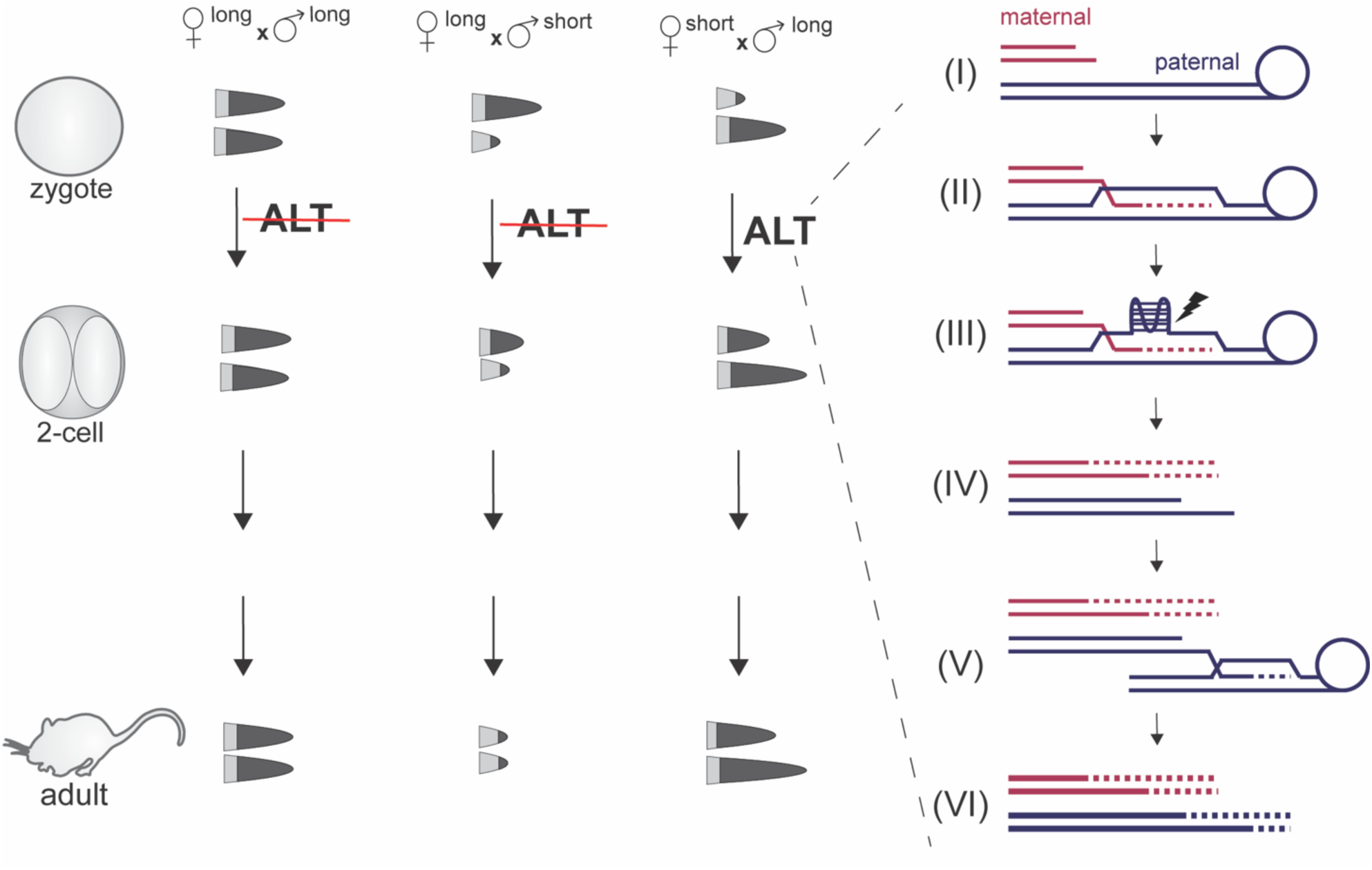
Model for a parent-of-origin effect on telomere elongation by ALT. Short maternal telomeres with single-stranded 3’ G-rich overhangs (I) initiate ALT by invading long paternal telomeres, leading to elongation of the G-rich maternal strand using the C-rich paternal strand as a template (II). The long G-rich paternal strand adopts a G4 structure that induces DNA damage (III). The DNA damage response generates a single-stranded paternal overhang, while the C-rich maternal strand elongates using the G-rich maternal strand as a template (IV). The paternal overhang invades another long paternal telomere, leading to elongation of the overhang (V) and, subsequently, the sister strand (VI).

This model requires the presence of a short telomere to initiate ALT and a long telomere to serve as the target of invasion and template for elongation. However, since reciprocal crosses are not equivalent, this elongation must also depend on an epigenetic difference between maternal and paternal telomeres. Maternal and paternal chromosomes at fertilization differ radically in chromatin state. The sperm-deposited paternal chromosomes initially lack H3K9me3 heterochromatin, and the paternal genome is rapidly demethylated after fertilization.^33^ In contrast, both H3K9me3 and methylated DNA are present on maternal chromosomes. Paternal chromosomes also lack the chromatin remodeling protein ATRX but are enriched for the histone chaperone DAXX compared to maternal chromosomes.^34–36^ We suspect that parental epigenetic asymmetry interacts with telomere length asymmetry to promote ALT. For example, epigenetic factors might destabilize T-loops on short maternal telomeres or stabilize T-loops on short paternal telomeres, making ALT dependent on both short maternal telomeres for single-stranded overhangs and long paternal telomeres as targets for invasion. The detailed mechanisms leading to ALT, or to telomere shortening in the reciprocal cross, are important avenues for future research.

## Methods

### Mice

*Mus musculus* mouse strains were purchased from the Jackson Laboratory: 129X1/SvJ, stock #000691; FVB/NJ, stock# 001800. *Mus spretus* was purchased from RIKEN BioResource Research Center (SPR2, RBRC00208). All mice used in this study were 7-8 weeks old females and 12-20 weeks old males. All animal experiments were approved by the Institutional Animal Care and Use Committee of the University of Pennsylvania and were consistent with the National Institutes of Health guidelines (protocol: #804882).

### Embryos and MII oocyte collection and culture

Female mice were super-ovulated by injecting 10 IU of pregnant mare serum gonadotropin (PMSG; Peptides International) followed by human chorionic gonadotropin (hCG; Sigma), either 56 hours after PMSG injection for the 129×1/SvJ strain or 48 hours after PMSG for the FVB/NJ strain. MII oocytes were collected 14-15 hours after hCG injection. Embryos were collected 18- 19 hours post-hCG after mating with male mice. MII oocytes and embryos were collected into M2 medium (Sigma; MR-015) and subsequently denuded of cumulus cells in M2 medium with hyaluronidase (0.15 mg/ml, Sigma, H3884). Embryos were cultured in EmbryoMax Advanced KSOM (AKSOM, Millipore Sigma, MR-101-D) in a humidified atmosphere of 5% CO_2_ in air at 37 °C. Embryos were cultured for 17 hours in AKSOM supplemented with the kinesin-5 inhibitor STLC (S-trityl-L-cysteine, Sigma, 10 μM) and the APC/C inhibitor proTAME (Medchemexpress, 5 μM) either 4 or 24 hours after collection for arrest at the one- or two-cell mitosis stage, respectively. To measure changes in telomere length between the first and second mitosis, mice for one-cell stage embryos were injected with PMSG 24 hours after mice for two-cell stage embryos. One-cell and two-cell stage embryos were then fixed simultaneously to ensure identical conditions for telomere FISH and immunostaining. G1 and S-G2 two-cell embryos were fixed 17-18 and 27-30 hours after embryo collection, respectively.^37^ EdU (5- ethynyl-2′-deoxyuridine) staining confirmed that embryos are in S-phase 28 hours after embryo collection (Figure S3). For arrest in mitosis at the morula to blastocyst stage, embryos were treated with STLC and proTAME for 4 hours, 72 hours after collection.

### Immunostaining

Embryos were exposed to acidic Tyrode’s solution (Sigma, T1788) for 2 minutes to remove the zona pellucida. After a brief recovery in fresh M2 medium, the embryos were fixed on slides in a humid chamber overnight with 1% paraformaldehyde in distilled water (pH 9.2) containing 0.15% Triton X-100 and 3 mM dithiothreitol. The slides were washed with phosphate buffered saline (PBS, Corning, 21-031-CV) three times, incubated in 0.5% Triton X-100 in PBS for 30 minutes, blocked in 3% BSA in PBS for 1 hour at room temperature, incubated overnight at 4°C or 1 hour at room temperature with primary antibodies, washed three times with 0.1% Tween 20 in PBS for 10 minutes each time, incubated with secondary antibodies for 1 hour at room temperature, washed three times again, and mounted in Vectashield with 4’,6-diamidino-2- phenylindole (DAPI, Vector) to visualize the chromosomes. To visualize G-quadruplex (G4) structures, slides were treated with a FLAG-tagged antibody that specifically binds to the G4 structure, followed by an antibody that recognizes the FLAG epitope tag, and finally, a fluorescence-labeled anti-FLAG antibody. Slides were imaged using a Leica TCS SP8 Four Channel Spectral Confocal microscope.

Primary antibodies used for immunostaining were anti-5hmC (Active motif, #39792), anti-TRF1 (Abcam, ab192629), anti-γH2AX (Abcam, ab81299), anti-DNA G-quadruplex structure (Millipore Sigma, MABE917), anti-FLAG (Cell signaling, #2368). Secondary antibodies were Alexa Fluor 488-conjugated anti-mouse (Invitrogen, A-21202), Alexa Fluor 488-conjugated anti-Rat (Invitrogen, A-21208), Alexa Fluor 647-conjugated anti-Rat (Invitrogen, A-21244) and Alexa Fluor 594-conjugated anti-rabbit (Invitrogen, A-21207).

### Fluorescence in situ Hybridization (FISH)

Metaphase II oocytes or embryos were exposed to a hypotonic solution (1% sodium citrate and 0.5% BSA) for 5 minutes, followed by fixation using methanol: acetic acid (3:1) on the slides. Slides were air-dried for 1 hour. Telomeres were denatured at room temperature for 10 minutes with 4N HCl and then neutralized with 100 mM Tris-HCl (pH 8.5) for 20 minutes. Telomeres were then hybridized with Alexa Fluor 488-labeled Tel-C probes (PNA Bio, F1004) at 500 nM. Slides were washed two times with 0.1% tween 20 in 2x Saline-Sodium Citrate buffer (SSC, Corning 46-020-CM) for 15 minutes each and mounted in Vectashield with DAPI.

For FISH combined with immunostaining of 5-hmC, slides were incubated in 3% BSA in PBS after the neutralization step, then incubated with the 5-hmC primary antibody for 1 hour at room temperature, washed three times (10 minutes each) with 0.1% Tween 20 in BSA buffer, incubated with secondary antibodies for 2 hours at room temperature, and washed three times again. Telomere hybridization, washing, and mounting was then performed as above.

For Native FISH, embryos were fixed on slides as for immunostaining. The slides were washed with PBS three times, incubated in 0.5% Triton X-100 in PBS for 30 minutes, then in 3% blocking reagent (Roche, #11096176001) in maleic acid buffer (100 mM maleic acid and 150 mM NaCl) supplemented with 500 μg/mL RNase A for 45 minutes at 37°C. After washing the slides three times with PBS at room temperature, embryos were dehydrated by consecutively incubating the slides in ethanol solutions at increasing concentrations (70%, 95%, and 100%) for 5 minutes each. Slides were air-dried at room temperature for two minutes to completely remove the 100% ethanol. Telomeres were then hybridized with Cy3-labeled Tel-G probes (PNA Bio, F1006) or Alexa Fluor 488-labeled Tel-C probes at 500 nM (PNA Bio, F1004) in the dark for 45 minutes at room temperature, washed twice with 70% formamide and 10 mM Tris-HCl in distilled water (pH 7.5) for 15 minutes each time, washed three times with PBS at room temperature, and mounted in Vectashield with DAPI.

#### EdU staining

Embryos for EdU labelling were collected 21, 28 and 32 hours after embryo collection. Embryos were cultured for 2 hours in AKSOM supplemented with the EdU. EdU was labeled using the Click-iT EdU Cell Proliferation Kit (Thermo Scientific, C10337) following the manufacture’s instruction.

### Image analysis

All image analysis was carried out using ImageJ/Fiji. To quantify telomere FISH signal intensities, circles of constant diameter were drawn around telomeres, and the average intensity was calculated for each telomere after subtracting the background obtained from nearby regions. Each data point represents a telomere of an individual chromatid in MII eggs or a pair of telomeres of sister chromatids in embryos, where individual chromatids could not be resolved reliably. For the inter-species crosses (Figure 1B), telomere intensities were normalized to the Mus musculus^long^♀ x Mus spretus^short^♂ cross for each experiment. For the intra-subspecies crosses (Figure 1C and 2A), telomere intensities were normalized to the 129^short^♀ x FVB^long^♂ cross for each experiment. For telomere elongation measurements (Figure 2B), telomere intensities were normalized to the maternal telomere length in one-cell 129^short^♀ x 129^short^♂ embryos for each experiment.

To quantify TTAGGG or AATCCC Native FISH signals in interphase nuclei (Figure 3B and 4A), a threshold intensity was set to differentiate FISH signal from background and then the number of regions was counted. The same threshold was used for embryos from each cross.

To measure G4 or γH2AX intensity at telomere regions, a threshold was set to define regions containing TRF1 signal, and the average intensity within each region was measured. The average background was then subtracted from these values. G4 or γH2AX intensities were summed for each cell.

### Statistical Analysis

Statistical analysis was performed with GraphPad Prism (GraphPad Software Inc.). Differences between groups were analyzed by Student’s t-test, and comparisons between more than two groups were analyzed by one-way ANOVA with Tukey’s multiple comparisons test. P < 0.05 was considered statistically significant.

## Supplementary Figures

**Figure S1.**
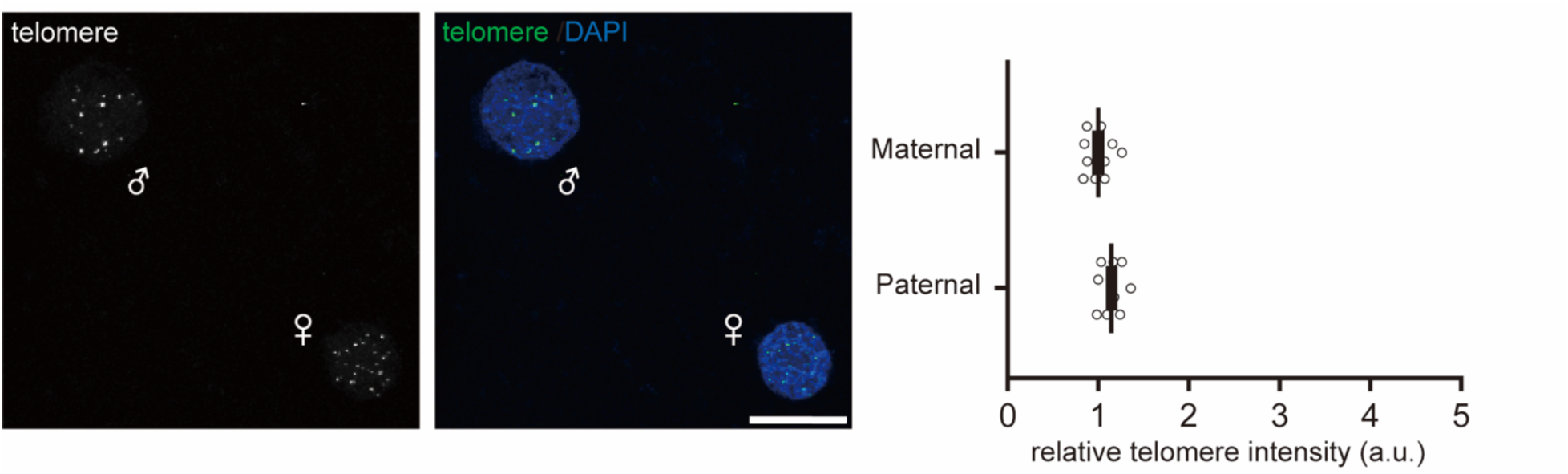
Telomere lengths of maternal and paternal pronuclei are similar. In contrast to previous reports^38–40^, we detected minimal differences in the telomere length transmitted by eggs and sperm from the same inbred strain, based on analysis of the female and male pronuclei in the zygote immediately following fertilization. Representative images show telomere FISH of zygotes from pure FVB^long^ strain; scale bar, 20 µm. Graph shows relative telomere intensities of maternal and paternal pronuclei. Each dot represents the total telomere intensity integrated over a single pronucleus; n=10 zygotes from 2 females; bars: mean and standard error of the mean. Data were normalized to the mean of the maternal pronuclei.

**Figure S2.**
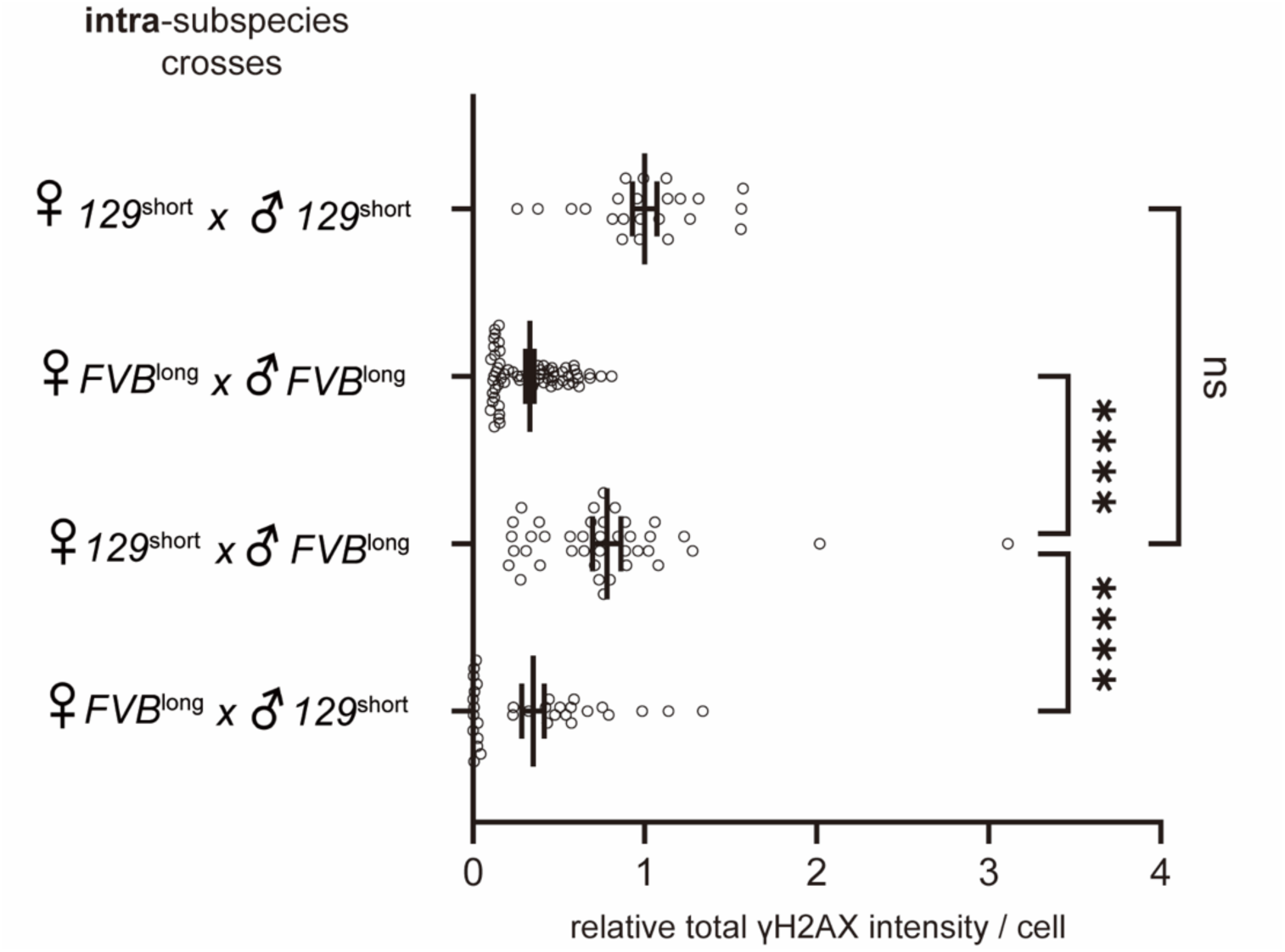
γH2AX staining in two-cell embryos. Analysis of total γH2AX signals for the data shown in Figure 4C. Data are presented as in Figure 4C, except that each dot represents γH2AX signal integrated over the entire nucleus rather than only overlapping telomeric foci.

**Figure S3.**
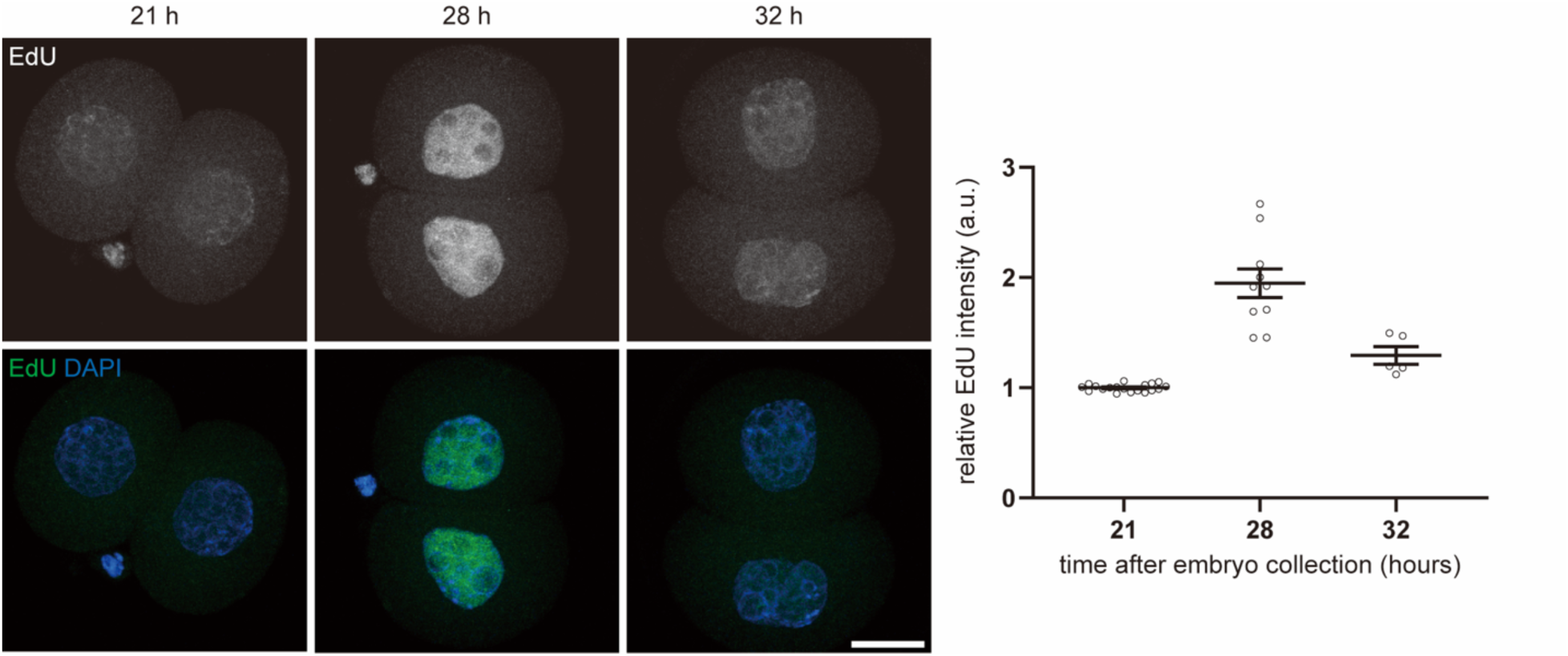
Timing of S phase in two-cell embryos. EdU staining of two-cell embryos from pure FVB^long^ strain. Representative images are shown; scale bar, 20 µm. Graph shows relative EdU intensities of the nuclei. Each dot represents the total EdU signal over one nucleus, normalized to the mean of the 21 h timepoint. For each sample, n ≥ 5 embryos were measured. Bars: standard error of the mean.

## Acknowledgments

We thank members of the Philadelphia Chromosome Club, Levine Lab, and Lampson Lab for helpful discussions. We also thank the Greenberg Lab for reagents and their vital input, especially that of T. Zhang, H. Jiang, and R. Greenberg. T. Zhang also provided valuable feedback on the manuscript. Funding was provided by NIH grants GM122475 (M.A.L.) and GM124684 (M.T.L), as well as seed grants from the Penn Center for Genome Integrity (M.A.L. and M.T.L.) and the University Research Fund from the University of Pennsylvania (M.A.L). H.J. was supported in part by a Postdoctoral Fellowship Program from the National Research Foundation of Korea.

